# The influenza A virus host shutoff factor PA-X is rapidly turned over in a strain-specific manner

**DOI:** 10.1101/2020.08.27.271320

**Authors:** Rachel Emily Levene, Shailab D. Shrestha, Marta Maria Gaglia

**Affiliations:** Graduate Program in Molecular Microbiology, Tufts University Graduate School of Biomedical Sciences, Boston, MA, USA; Department of Molecular Biology and Microbiology, Tufts University School of Medicine, Boston, MA, USA

**Keywords:** influenza A virus, PA-X, host shutoff, protein turnover

## Abstract

The influenza A endoribonuclease PA-X regulates virulence and transmission of the virus by reducing host gene expression and thus regulating immune responses to influenza A virus. Despite this key function in viral biology, the levels of PA-X protein remain markedly low during infection, and previous results suggest that these low levels are not solely the result of regulation of the level of translation and RNA stability. How PA-X is regulated post-translationally remains unknown. We now report that the PA-X protein is rapidly turned over. PA-X from multiple viral strains are short-lived, although the half-life of PA-X ranges from ∼30 minutes to ∼3.5 hours depending on the strain. Moreover, sequences in the variable PA-X C-terminal domain are primarily responsible for regulating PA-X half-life, although the N-terminal domain also accounts for some differences among strains. Interestingly, we find that the PA-X from the 2009 pandemic H1N1 strain has a longer half-life compared to the other variants we tested. This PA-X isoform has been reported to have a higher host shutoff activity, suggesting a role for protein turnover in regulating PA-X activity. Collectively, this study reveals a novel regulatory mechanism of PA-X protein levels that may impact host shutoff activity during influenza A virus infection.

**IMPORTANCE:** The PA-X protein from influenza A virus reduces host immune responses to infection through suppressing host gene expression, including genes encoding the antiviral response. Thus, it plays a central role in influenza A virus biology. Despite its key function, PA-X was only discovered in 2012 and much remains to be learned including how PA-X activity is regulated to promote optimal levels of viral infection. In this study, we reveal that PA-X protein levels are very low likely because of rapid turnover. We show that instability is a conserved property among PA-X variants from different strains of influenza A virus, but that the half-lives of PA-X variants differ. Moreover, the longer half-life of PA-X from the 2009 pandemic H1N1 strain correlates with its reported higher activity. Therefore, PA-X stability may be a way to regulate its activity and may contribute to the differential virulence of influenza A virus strains.

## INTRODUCTION

Viruses employ multiple strategies to block host antiviral defenses. In many viruses, this inhibition is partly achieved through a blockade of host gene expression, a process termed “host shutoff”. One of the ways influenza A virus induces host shutoff is through the activity of a viral endoribonuclease (RNase), PA-X (1–3). The RNase activity of PA-X results in global changes in the cellular transcriptome (3, 4), while viral mRNAs and genomic RNAs escape PA-X degradation (5). Influenza A viruses engineered to lack PA-X trigger a more potent innate immune and inflammatory response in mice, chickens, and pigs, presumably because of decreased host shutoff (1, 6–10). These results indicate that PA-X controls immune responses *in vivo*. More recently, a role for PA-X in transmission has also been established (11, 12). Despite its clear *in vivo* role, much remains to be determined about PA-X biology during infection. PA-X was discovered in 2012 and studies to date have primarily focused on dissecting its mechanism of action, including how subsets of RNAs, including viral RNAs, escape PA-X degradation. Whether and how PA-X activity itself is regulated has not been explored in detail. One study found that co-translational N-terminal acetylation is required for PA-X host shutoff activity (13), but whether there are other co- and post-translational regulatory mechanisms remains unknown. Interestingly, prolonged PA-X expression may be cytotoxic, as shown in yeast (13, 14). While this is not surprising given its widespread downregulation of RNAs, this observation underscores the idea that PA-X activity may need to be regulated during infection to prevent premature death of the infected cells.

One way to regulate or limit PA-X activity could be to keep pools of active PA-X protein low in the infected cell. Indeed, PA-X protein levels are remarkably low during infection. Metabolic labeling of nascent proteins in influenza A virus infected cells showed that the levels of PA-X are barely above the limit of detection and lower than other influenza proteins, including the well-known host shutoff factor NS1 (1). These low levels are in part due to how PA-X is produced. PA-X production requires a programmed +1 ribosomal frameshift during translation of segment 3, which encodes the polymerase acidic (PA) subunit of the viral RNA dependent RNA polymerase (RdRp) (1, 15). Only a small percentage of translational runs result in frameshifting and produce PA-X, which has the same N-terminal RNase domain as PA (PA-X amino acids (aa) 1-191) and a unique C-terminal domain known as the “X-ORF” (PA-X aa 192-232 or 192-252 depending on the strain) (1, 16). Of note, despite this non-canonical production mechanism, PA-X is present in all influenza A virus isolates (16), which underscores its importance in influenza A virus biology. However, the mechanism of PA-X production does not entirely explain the low levels of PA-X. Indeed, PA-X levels are also low or undetectable when PA-X is expressed ectopically (17). In these experiments, a single nucleotide has been deleted in the expression construct to mimic the frameshift and all translational runs produce PA-X (17). The low levels are also not because PA-X degrades its own mRNA, as mutating the PA-X catalytic residues D108 or K134 in this ectopic expression system does not increase PA-X levels dramatically (3, 17). This suggests that PA-X levels may be regulated at the level of protein stability.

Regulation of stability is a well-established post-translational mechanism of protein regulation. For example, Kosik *et. al* showed that protein turnover regulates the immunomodulatory activity of PB1-F2, another influenza A virus virulence factor (18). PB1-F2 inhibits induction of the antiviral cytokine interferon beta (IFN-β) and increases the activity of the viral RdRp (19–21). Mutating the residues required for PB1-F2 ubiquitination and rapid turnover increases PB1-F2 protein levels, which results in a stronger inhibition of IFN-β induction (18). Similarly, regulation of PA-X at the level of protein stability could modulate the host shutoff activity of influenza A virus.

Previous observations suggest that the X-ORF may regulate the levels of the PA-X protein, as ectopic expression of the N-terminal domain of PA-X results in much higher protein levels than ectopic expression of either wild type (wt) or catalytically inactive PA-X (17). This is not due to changes in frameshifting, as this ectopic expression construct has a nucleotide deletion to mimic the frameshift and only encodes PA-X. A role for the X-ORF in post-translational control of PA-X levels and activity would explain why almost all influenza A strains encode a PA-X protein that has a 61 or 41-aa X-ORF (16) even though only the first 15 aa of the X-ORF are required for PA-X shutoff activity in cells (3, 17, 22). Moreover, most of the sequence variation among PA-X from different strains occurs after the 15th amino acid of the X-ORF, and the strength of PA-X activity has been reported to vary among strains (17, 23, 24).

Here we report the first investigation of post-translational regulation of PA-X protein levels. We find that PA-X is markedly unstable and identify a region of the A/Puerto Rico/8/1934 H1N1 (PR8) X-ORF that regulates PA-X turnover. Additionally, we find that the half-life of PA-X differs among strains, even though all the PA-X variants tested have generally short half-lives. Interestingly, of the variants we tested, PA-X from the 2009 pandemic H1N1 strain had the longest half-life, which may explain its reported higher host shutoff activity (17, 23, 24). Thus, modulation of PA-X stability may be a mechanism used to regulate PA-X host shutoff activity in different strains and may contribute to influenza A pathogenesis.

## RESULTS

### PA-X is unstable

Several observations suggested to us that PA-X may be unstable. PA-X is expressed at low levels not only during infection (1), but also during ectopic overexpression from a high expression CMV promoter [(17), our unpublished observations]. Furthermore, ectopic overexpression of the N-terminal RNase domain from the same high expression CMV promoter results in much higher protein levels (17). While the RNase domain in isolation is inactive in cells (3, 5, 17, 22), the differences in protein levels are not due to the loss of activity, because expressing a catalytically inactive version of PA-X still results in markedly low PA-X expression levels (3, 17). Thus, we sought to directly test if PA-X is rapidly turned over. To determine the half-life of the PA-X protein, we performed a cycloheximide chase in human embryonic kidney (HEK) 293T cells transfected with a C-terminally myc tagged PA-X from the influenza A virus strain A/Puerto Rico/8/1934 H1N1 (PR8) containing the D108A mutation in the RNase active site. This mutation abolishes PA-X catalytic activity and host shutoff function (3, 5). We used catalytically inactive PA-X for this experiment and throughout this study to eliminate any confound from changes in host gene expression or PA-X mRNA levels as a result of PA-X activity. Additionally, to make sure any differences in PA-X protein levels are due to changes in turnover and not frameshifting, in this experiment and throughout this study we used PA-X expression constructs where all translational runs produce PA-X, as a single nucleotide was deleted to mimic the frameshift. After treating PA-X-expressing cells with cycloheximide to inhibit translation, we analyzed PA-X protein levels at 0, 15, 30, 60, and 120 minutes after cycloheximide addition by western blotting. Strikingly, PA-X protein levels decreased dramatically one hour after the addition of cycloheximide (Fig 1A and 1B), suggesting that PA-X is markedly unstable. By contrast, levels of PA-X mRNA did not decrease throughout the cycloheximide time course (Fig 1C), confirming that the decrease in PA-X protein levels is not due to changes in transcript levels. Based on regression analysis, we calculated a half-life of 86 minutes for PR8 PA-X (Fig 1B, Table 1). This is short compared to the half-life of other influenza proteins. For example, the immune evasion factor NS1 from PR8 has a half-life of over 24 hours (25) and PR8 PB1-F2 has a half-life of 6.5 hours (26). In addition, human proteins have a median half-life of 8.7 hours (27). Together, these data show that PA-X is rapidly turned over compared to other viral and host proteins.

**TABLE 1:** HALF-LIVES OF PA-X FROM PR8, pH1N1, UDORN, HK06, AND PERTH. A linear regression was carried out on the quantification plots in Figure 3. The slope of the linear regression line and the calculated half-life are reported. The half-life was calculated as the negative of the inverse of the slope in minutes. The slope and half-life for PR8 PA-X was calculated based on regression analysis from Figure 1B.

| PA-X | Slope of linear regression | Half-Life in Minutes | 95% confidence interval on slope |
| --- | --- | --- | --- |
| A/Puerto Rico/8/1934 | -0.70 | 86 | -0.90 to -0.50 |
| A/California/7/2009pdm pH1N1 | -0.29 | 207 | -0.36 to -0.22 |
| A/Udorn/307/1973 H3N2 | -1.77 | 34 | -2.41 to -1.14 |
| A/Perth/16/2009 H3N2 | -0.39 | 156 | -0.48 to -0.29 |
| A/Hong Kong/218847/2006 H1N1 | -0.73 | 83 | -1.07 to -0.38 |

**FIGURE 1:**
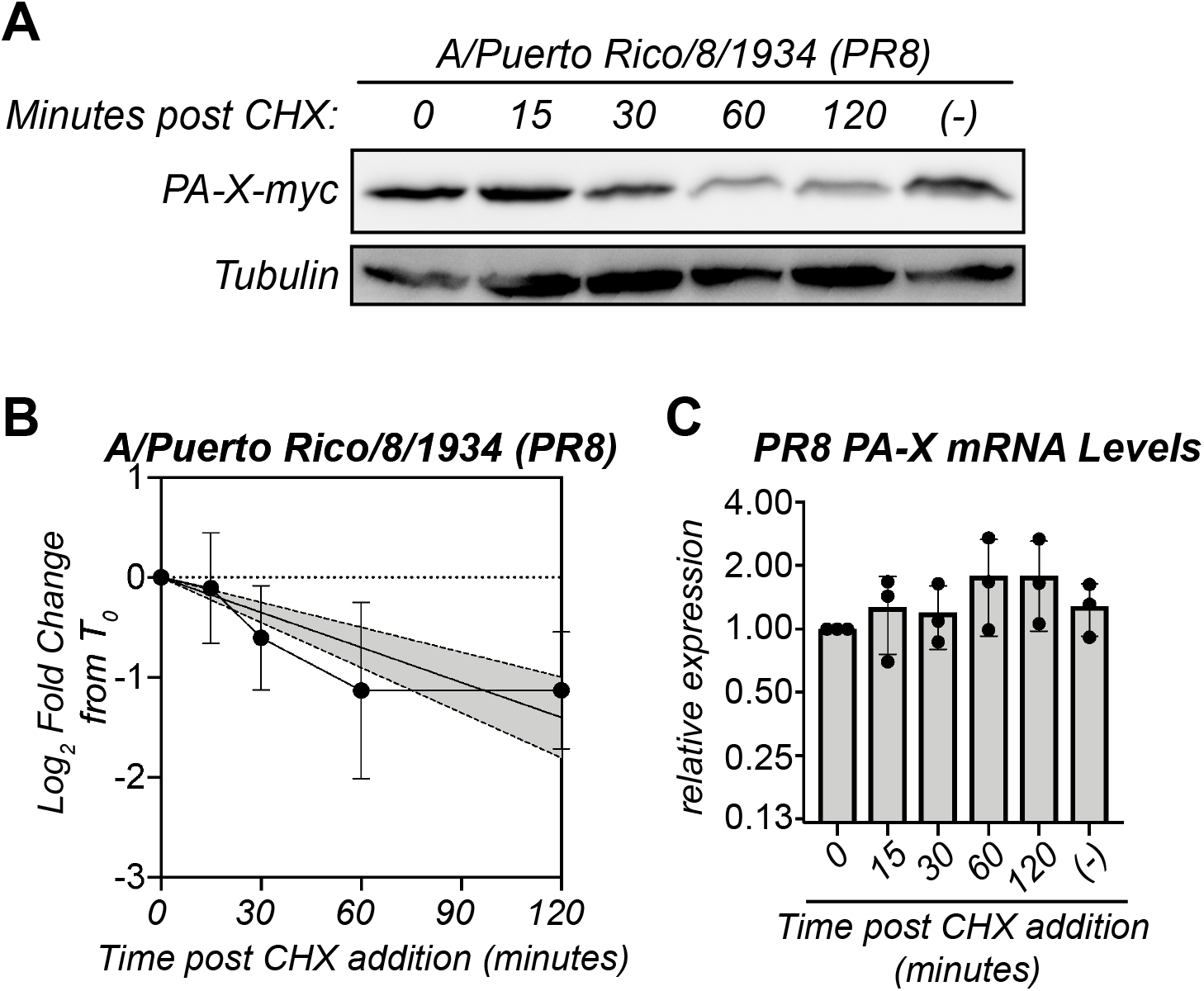
PA-X is unstable. (A-C) Protein and RNA samples were collected from HEK293T cells expressing catalytically inactive PR8 PA-X D108A at the indicated time points after treatment with 200 μg/mL of cycloheximide (CHX) to inhibit translation. (−) indicates vehicle-treated cells collected at 120 minutes. (A) Representative western blot using anti-myc antibodies to detect myc-tagged PA-X and tubulin as a loading control. (B) Plot of the relative levels of PA-X after cycloheximide treatment. For each replicate, PA-X protein levels were normalized to tubulin protein levels and reported as the log_2_ transformed ratio relative to time 0. The line represents the linear regression and the shaded region its 95% confidence interval. n = 7 (C) Levels of PA-X mRNA were measured by RT-qPCR. After normalization to 18S rRNA, levels were plotted as amounts relative to time 0. n = 3.

### PA-X host shutoff activity in infected cells is reduced upon translational inhibition

To confirm that PA-X is unstable in the context of influenza A virus infection we first attempted to measure its half-life in PR8-infected cells using an antibody against PR8 PA that has been previously used to detect PR8 PA-X (28). However, like Rigby *et al*. (28), we observed multiple bands in the section of the blot corresponding to the predicted molecular weight of PA-X, and were unable to reliably and reproducibly identify the band corresponding to PA-X. Unfortunately, we cannot tag PA-X in the context of the virus. The C-terminus cannot be tagged because it overlaps with the coding sequence for the C-terminus of PA (1, 15). N-terminal tags reduce PA activity and viral replication (29, 30), and are also expected to reduce the activity of PA-X because of the importance of its N-terminal acetylation (13). Therefore, we sought another way to look for evidence of PA-X instability during infection and used PA-X activity in the presence of cycloheximide as a proxy for rapid changes in PA-X protein levels. We reasoned that if levels of PA-X are quickly depleted after halting translation with cycloheximide, there will be less PA-X host shutoff activity in cells. To confirm this, we first tested PA-X host shutoff activity in the presence of cycloheximide in A549 cells that inducibly express either wt PR8 PA-X or the catalytically inactive PR8 PA-X D108A mutant. These cells allow us to reliably measure PA-X-dependent changes in the RNA levels of endogenous genes (3, 5). Following overnight induction of PA-X expression and a subsequent 2-hour cycloheximide treatment, we analyzed levels of G6PD and TAF7 mRNA, which are representative PA-X target and resistant RNAs, respectively (3). Levels of G6PD significantly decreased upon PA-X expression in vehicle-treated cells (Fig 2A). In contrast, there was no statistically significant decrease in G6PD levels in PA-X-expressing cells treated with cycloheximide (Fig 2A). We note that there was substantial experimental variability in cycloheximide-treated cells, perhaps due to different levels of PA-X mRNA or of residual PA-X protein after treatment (Fig 2A). Nonetheless, cycloheximide had no effect on the levels of G6PD in cells expressing catalytically inactive PA-X or levels of the PA-X resistant RNA TAF7 (Fig 2A), confirming that the effect of cycloheximide on RNA levels was at least partially linked to PA-X activity. Furthermore, PA-X mRNA levels did not decrease with cycloheximide (Fig 2A), confirming that any decrease in PA-X host shutoff activity is not the result of a decrease in PA-X transcript levels.

**FIGURE 2:**
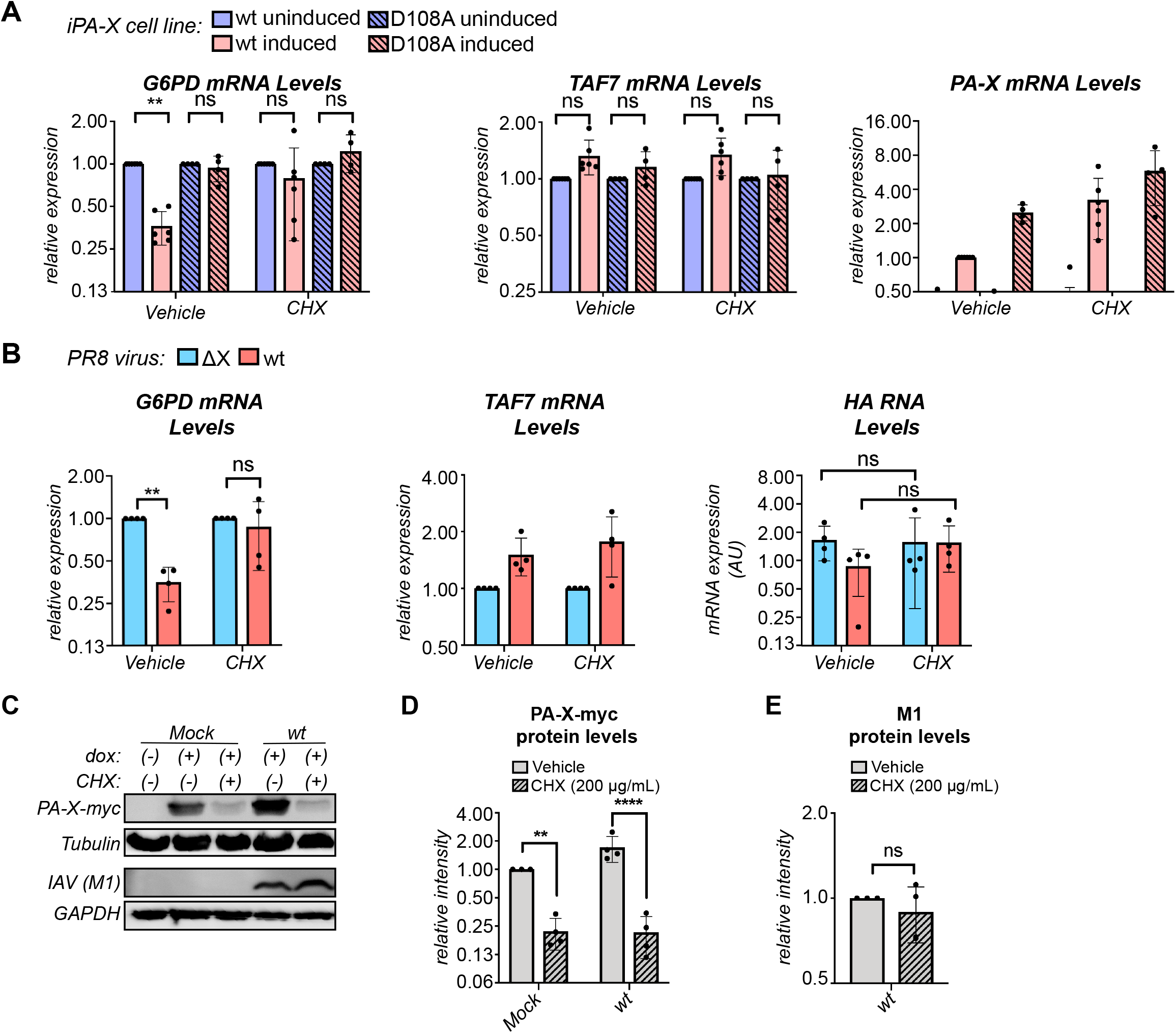
PA-X is unstable during infection. (A) Levels of the indicated endogenous RNAs were measured by RT-qPCR in A549 “iPA-X” cells, which express wild-type (wt) PR8 PA-X or a PR8 PA-X catalytic mutant (D108A) under a doxycycline-inducible promoter. RNA samples were collected after 14 hours of PA-X induction with doxycycline followed by a 2-hour treatment with 200 μg/mL of cycloheximide (CHX) or vehicle control. After normalization to 18S rRNA, mRNA levels are plotted relative to uninduced cells for each condition and cell line (G6PD, TAF7) or relative to induced wt iPA-X cells (PA-X). n ≥ 4. (B) Levels of the endogenous human mRNAs (G6PD and TAF7) and viral RNAs (HA) were measured by RT-qPCR in cells infected with wt PR8 or the PA-X-defective PR8 ΔX strain at a MOI of 5 for 24 hours prior to a 2-hour treatment with 200 μg/mL of cycloheximide (CHX) to inhibit translation or a vehicle control. After normalization to 18S rRNA, RNA levels are plotted relative to PR8 ΔX-infected cells for each condition (G6PD, TAF7) or as amounts normalized to cellular 18S rRNA (HA). n = 4. (C-E) Expression of PA-X-myc was induced in HEK293T-iPA-X_PR8_D108A with 2.0 μg/mL doxycycline (dox) for 6 hours, followed by either mock or wt PR8 infection at MOI = 1 for 18 hours. Protein samples were collected after a subsequent 2-hour treatment with 200 μg/mL of cycloheximide (CHX) to inhibit translation (+) or with vehicle control (−). (C) Representative western blot using anti-myc antibodies to detect myc-tagged PA-X with tubulin as a loading control and anti-IAV antibodies to detect M1 with GAPDH as a loading control. (D) For each replicate, levels of myc-tagged PA-X were normalized to tubulin protein levels and reported as the amount relative to induced, mock infected, vehicle treated cells. n = 4. (E) For each replicate, levels of M1 were normalized to GAPDH protein levels and reported as the amount relative to wt infected, vehicle treated cells. N = 4. ** = *p* < 0.01, **** = *p* < 0.0001, ns = *p* > 0.05. Significance was calculated using 3-way ANOVA followed by Tukey’s multiple comparisons test (A), 2-way ANOVA followed by Sidak’s multiple comparisons test (B, D), or Student’s t-test (E).

We then performed the same experiment in A549 cells infected with either wt PR8 or PA-X-deficient PR8 virus (ΔX). PR8 ΔX carries mutations in the frameshifting site in segment 3 that reduce PA-X production, as well as a nonsense mutation at PA-X aa 201, which abolishes activity of any residual PA-X (3). At 24 hours post-infection, we analyzed levels of G6PD and TAF7 mRNA after a 2-hour cycloheximide treatment. As expected, G6PD levels were significantly lower in wt PR8-infected cells compared to PR8 ΔX-infected cells due to PA-X mediated host shutoff, while the levels of the PA-X-resistant RNA TAF7 were similar (Fig 2B). However, there was no significant difference in G6PD levels in wt PR8 *vs*. PR8 ΔX-infected cells after treatment with cycloheximide (Fig 2B). This result suggests that cycloheximide treatment prevents PA-X-mediated host shutoff, presumably due to a rapid decrease in PA-X protein levels. As expected, PA-X had no effect on TAF7 levels (Fig 2B). There were no significant changes in the RNA levels of the viral gene HA during cycloheximide treatment (Fig 2B), confirming that the decrease in PA-X activity is not the result of changes in viral replication. Thus, while we cannot examine changes in PA-X protein levels in infected cells and test its protein stability, we do observe markedly reduced PA-X activity after translation is inhibited. Taken together with the short half-life of PR8 PA-X (Fig 1) and the decrease in its activity after translational block during ectopic expression (Fig 2A), our results indicate that the reduced PA-X activity after translational block during infection is likely the result of decreased PA-X protein levels. Therefore, it is likely that PA-X is rapidly turned over during infection.

### PA-X-myc is unstable in influenza A virus-infected cells

Because we were unable to reliably detect PA-X during infection and there was some variability in measuring PA-X activity as a proxy for rapid changes in PA-X protein levels (Fig 2A), we sought an additional way to look for evidence of PA-X instability during infection. Thus, we measured the stability of ectopically expressed PA-X in mock or wt PR8-infected cells. For this experiment, we used HEK 293T cells that inducibly express C-terminally myc tagged catalytically inactive PR8 PA-X (HEK 293T-iPA-X_PR8_D108A), infected them for 18 hours and then treated them with cycloheximide to inhibit translation. As expected based on our previous results, PA-X-myc protein levels significantly decreased in both mock and wt PR8-infected cells following a 2-hour cycloheximide treatment (Fig 2C and 2D). In contrast, there were no significant changes in the levels of the virus-encoded M1 protein between vehicle- and cycloheximide-treated wt PR8-infected cells (Fig 2C and 2E), indicating that changes in PA-X-myc protein levels were not due to changes in viral replication. Thus, ectopically expressed PA-X-myc is rapidly turned over in infected cells, which is consistent with the idea that virus-encoded PA-X is also unstable under these conditions.

### The half-life of PA-X varies among influenza A virus strains

Although all influenza A virus strains are predicted to encode PA-X, the sequence of PA-X varies across strains (16). Thus, we sought to determine whether the instability we observed in PR8 PA-X was conserved in PA-X from other strains of influenza A virus. We created catalytically inactive D108A mutants of the PA-X from the 2009 pandemic H1N1 strain A/California/7/2009pdm (pH1N1), the lab-adapted H3N2 strain A/Udorn/307/1972 (Udorn), the seasonal H3N2 strain A/Perth/16/2009 (Perth), and the pre-pandemic seasonal H1N1 strain A/Hong Kong/218847/2006 (HK06), all with a C-terminal myc tag. Fusing all variants to the same tag allows us to better compare protein levels among strains, since many antibodies against influenza proteins only recognize a particular strain or serotype. Udorn and HK06 PA-X were as or more unstable than PR8 PA-X, with half-lives of 34 and 83 minutes, respectively (Fig 3A, 3B, 3D, 3E, Table 1). In contrast, Perth PA-X was more stable than PR8 PA-X, with a half-life of 156 minutes (Fig 3G, 3H, Table 1). pH1N1 PA-X was also significantly more stable than its PR8 counterpart, with a half-life of 207 minutes (Fig 3J, 3K, Table 1). Importantly, the longer lived PA-X variants (Perth and pH1N1 PA-X) still have a shorter half-life than the average human protein (∼9 hrs) (27), suggesting they are still relatively unstable. Also, there were no significant decreases in mRNA levels of PA-X from any of the strains during the cycloheximide time courses (Fig 3C, 3F, 3I, 3L), confirming that the decrease in PA-X protein levels is not due to changes in transcript levels. Together, these data show that instability is a conserved property among PA-X variants, but that there are differences in PA-X half-life among strains. Moreover, the pH1N1 PA-X is substantially longer lived than other variants.

**FIGURE 3:**
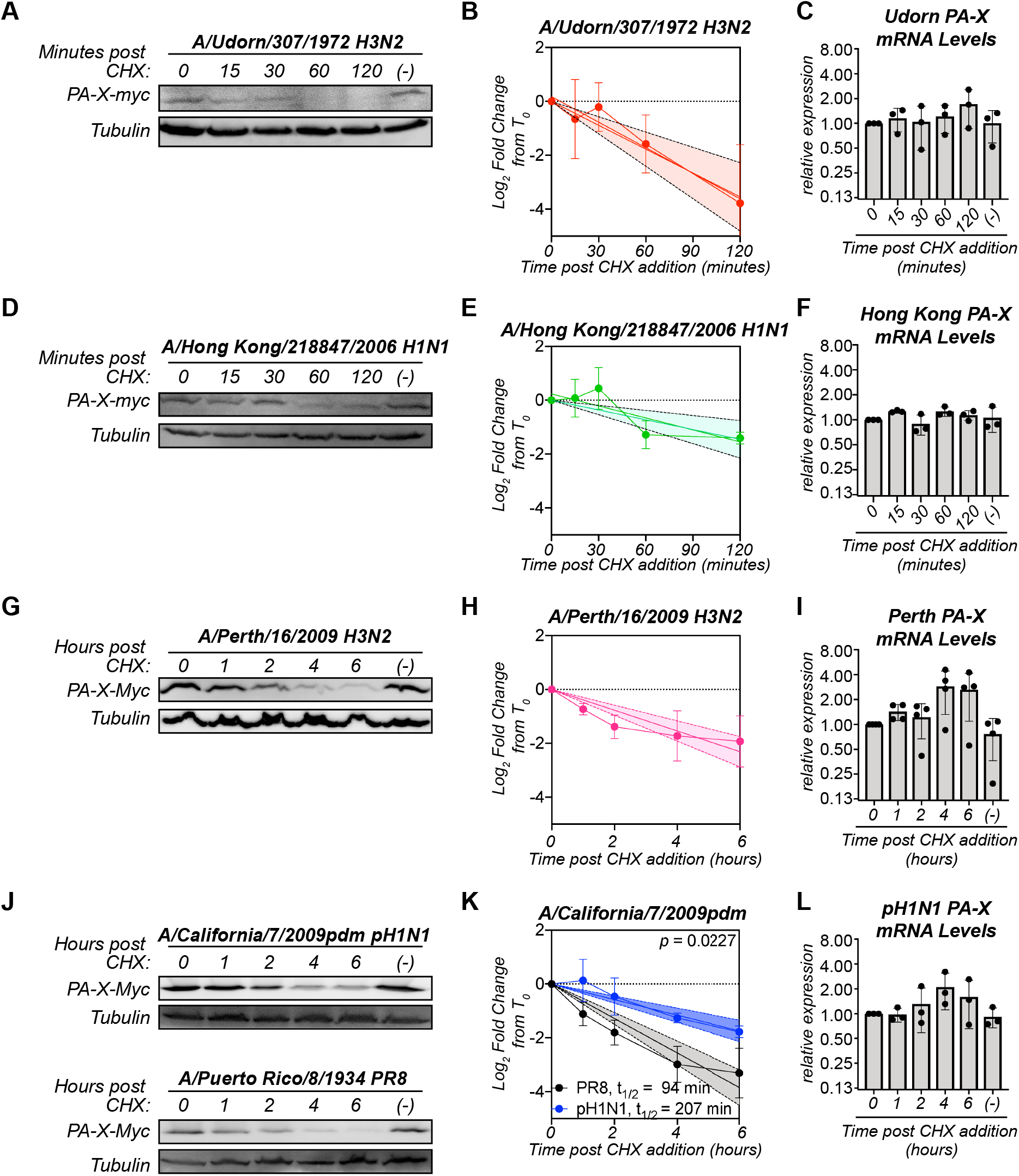
The half-life of PA-X varies among influenza A virus strains. Protein and RNA samples were collected from HEK293T cells expressing catalytically inactive (D108A mutant) C-terminally myc tagged PA-X from the indicated strains at the indicated time points after treatment with 200 μg/mL of cycloheximide (CHX) to inhibit translation. (−) denotes vehicle-treated cells collected at 120 minutes (A-F) or 6 hours (G-L). (A, D, G, J) Representative western blots using anti-myc antibodies to detect myc-tagged PA-X and tubulin as a loading control. (B, E, H, K) Plot of the relative levels of PA-X after cycloheximide treatment. For each replicate, PA-X protein levels were normalized to tubulin protein levels and reported as the log_2_ transformed ratio from time zero. The line represents the linear regression and the shaded region its 95% confidence interval. n ≥ 3 for each variant. The half-life (t_1/2_) is calculated as the negative inverse of the slope and reported in minutes in the figure and/or in Table 1. For (K), the *p* value is calculated relative to PR8 PA-X (PR8 PA-X: slope = −0.64, confidence interval: −0.75 – −0.53; pH1N1 PA-X: slope = −0.29, confidence interval: −0.36 – −0.22). (C, F, I, L) Levels of PA-X mRNA were measured by RT-qPCR. After normalization to 18S rRNA, levels are plotted as amounts relative to time 0. n ≥ 3.

### Amino acids 17-41 of the X-ORF contribute to regulation of PA-X instability

Because ectopic overexpression of the N-terminal domain of PA-X results in higher protein levels than overexpression of full length PA-X (17), we tested whether the X-ORF regulates PA-X stability. We took two complementary approaches: we fused the PR8 X-ORF to the stable reporter protein dsRed (“dsRed-XORF”) to test whether the X-ORF reduces DsRed stability and we truncated the X-ORF in PR8 PA-X to test whether this increases PA-X stability. We observed that levels of dsRed-XORF significantly decreased after six hours of cycloheximide treatment, whereas levels of dsRed remained constant (Fig 4A-B). There was no decrease in mRNA levels of dsRed or dsRed-XORF upon cycloheximide treatment (Fig 4C), confirming that decreases in dsRed protein levels are not due to changes in transcript levels. This indicates that addition of the X-ORF can reduce stability of an unrelated protein. To test the effect of X-ORF truncations, we chose to test a truncated version that ended after aa 16 of the X-ORF in PR8 PA-X (“PA-X16”, Fig 4D), because truncation of the entire X-ORF results in loss of nuclear localization of PA-X and host shutoff activity in cells (5, 17). In contrast, PA-X mutants retaining at least 15 aa of the X-ORF are primarily localized to the nucleus, like full-length PA-X (17) and have wild-type host shutoff activity (3, 17, 22). Thus, using the PA-X16 truncation should exclude potential confounding factors, such as changes in subcellular localization or in protein-protein interactions needed for activity. In contrast to the rapid decrease in the levels of full-length PA-X (PA-X61) after cycloheximide addition, the protein levels of PA-X16 remained constant throughout the two-hour time course (Fig 4E and 4F). There were also no significant changes in the mRNA levels of PA-X16 throughout the time course (Fig 4G). Thus, the truncated PA-X16 is more stable than the full-length protein. Together, these data show that the X-ORF, in particular sequences in the X-ORF after the 16^th^ residue, contains sequences that reduce protein stability and contribute to the instability of PA-X. Since the first 15 amino acids of the X-ORF are necessary and sufficient for host shutoff activity (3, 17, 22), these data also suggest that the portions of the X-ORF involved in mediating host shutoff activity and stability are different.

**FIGURE 4:**
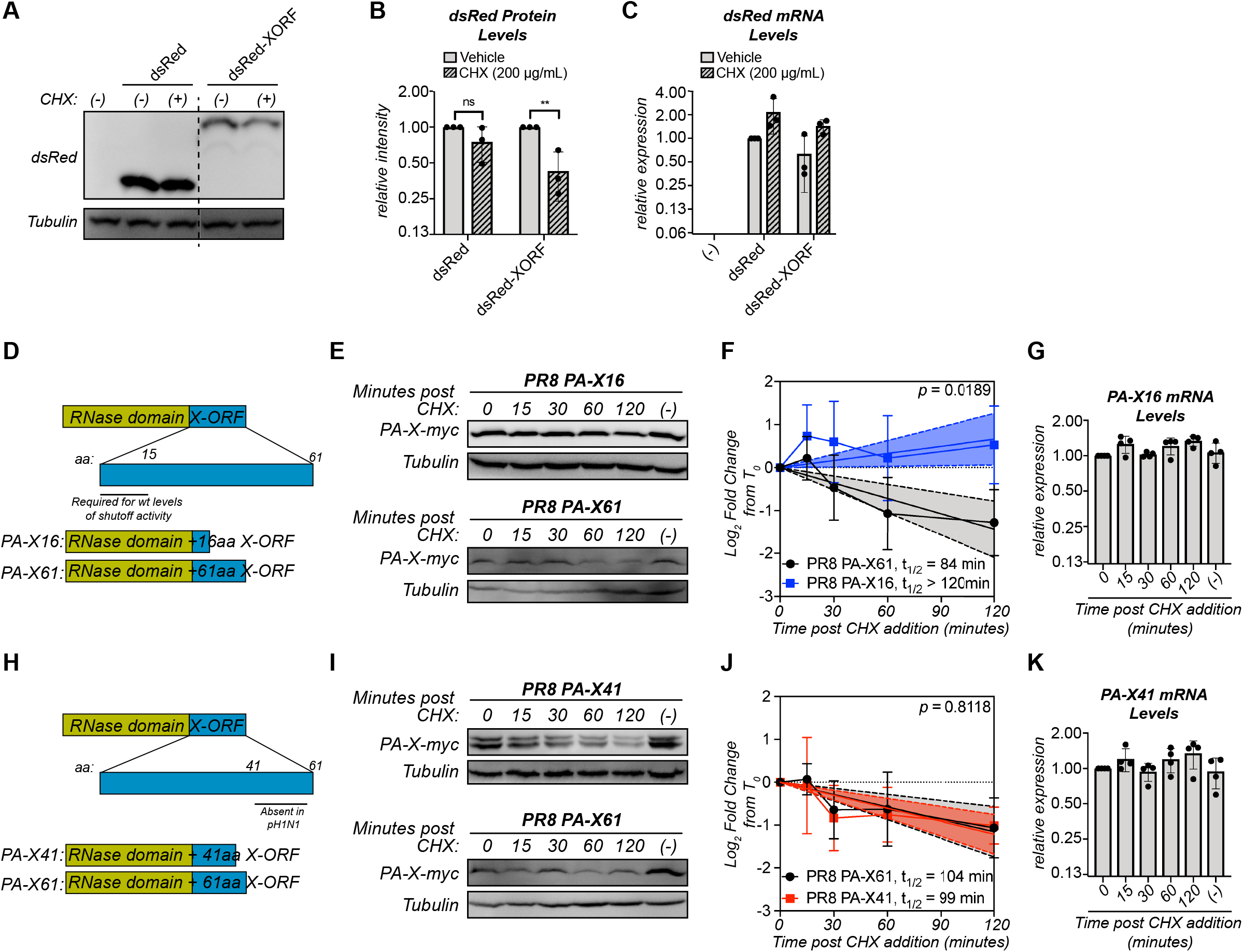
Amino acids 17-41 of the X-ORF contribute to regulation of PA-X turnover. (A-C) Protein and RNA samples were collected from HEK293T cells expressing the indicated dsRed variant 6 hours after treatment with 200 μg/mL of cycloheximide (CHX) to inhibit translation (+) or vehicle control (−). (A) Representative western blot using anti-dsRed antibodies to detect dsRed and tubulin as a loading control. (B) For each replicate, dsRed protein levels were normalized to tubulin protein levels and reported as the amount normalized to vehicle control for each dsRed variant. ** = *p* < 0.01, ns = *p* > 0.05, 2-way ANOVA followed by Sidak’s multiple comparisons test. n = 3. (C) Levels of dsRed mRNA were measured by RT-qPCR. After normalization to 18S rRNA, levels are plotted as amounts relative to dsRed vehicle control. n = 3. (D, H) Schematic diagram of PA-X domains and PR8 PA-X16 (D) or PR8 PA-X41 (H) truncations used. (E-G, I-K) Protein and RNA samples were collected from HEK293T cells expressing the catalytically inactive full-length or the indicated truncated C-terminally myc-tagged PA-X at the indicated time points after treatment with 200 μg/mL of cycloheximide (CHX) to inhibit translation. (−) indicates vehicle-treated cells collected at 120 minutes. (E, I) Representative western blot using anti-myc antibodies to detect myc-tagged PA-X and tubulin as a loading control. (F, J) Plot of the relative levels of PA-X after cycloheximide treatment. For each replicate, PA-X protein levels were normalized to tubulin protein levels and reported as the log_2_ transformed ratio from time 0. A linear regression was performed for each dataset and the *p* value calculated relative to PR8 PA-X61 (full length PA-X). The lines represent the linear regression and the shaded regions indicate the 95% confidence interval (F: PA-X61: slope = −0.72, confidence interval = −1.05 – −0.39; PA-X16: slope = 0.33, confidence interval = 0.03 – 0.63; J: PA-X61: slope = −0.58; confidence interval: −0.87 – −0.28; PA-X41: slope = −0.61, confidence interval = −0.84 – −0.38). The half-lives (t_1/2_) were calculated as the negative inverse of the slope and are reported in minutes. As the slope for PA-X16 is positive, the half-life of this protein cannot be calculated and is therefore reported as longer than the time course of the experiment. n ≥ 3 (G, K) Levels of the mRNAs for the indicated truncated PA-X were measured by RT-qPCR. After normalization to 18S rRNA, levels are plotted as amounts relative to time 0. n > 3.

While the first 15 aa of the X-ORF are generally conserved among strains, the sequences after the 16th aa vary. In particular, there are two main variants of PA-X in influenza A strains, which are distinguishable by their protein lengths: one is 252-aa long, with an X-ORF of 61 aa, and the other is 232-aa long, with an X-ORF of 41 aa (16). The PA-X variants in PR8, Perth, Udorn and HK06 all have a 61-aa X-ORF, like most human strains, while the pH1N1 PA-X has a 41-aa X-ORF, which is otherwise found in a subset of swine and canine strains (16). Although some studies have suggested that the length of the X-ORF may regulate activity (12, 31–33), it is still unclear exactly what the function of the additional 20 aa is. Since pH1N1 PA-X has a much longer half-life than most of the other PA-X isoforms we tested (Fig 3K, Table 1), we wondered if the shorter X-ORF accounted for the difference in stability. We thus measured the half-life of a truncated version of catalytically inactive PR8 PA-X that ended after the 41st amino acid with a C-terminal myc tag, which we termed “PA-X41” (Fig 4H). Despite the truncation, PR8 PA-X41 protein levels rapidly decreased after translation inhibition (Fig 4I and 4J), with a half-life of 99 minutes. This is not significantly different from that of full length PR8 PA-X (PA-X61) (*p* = 0.8118) (Fig 4J). We note that we consistently detected a double band in cells expressing PR8 PA-X41. While the reason for this doublet is currently unknown, a similar doublet was also apparent in the staining for pH1N1 PA-X in Hayashi *et al*. (17). There were no significant decreases in the mRNA levels of PR8 PA-X41 (Fig 4K), confirming that the decrease in PR8 PA-X41 protein levels is not due to changes in transcript levels. Thus, the distal 20 residues of PR8 PA-X likely do not play a role in PR8 PA-X’s rapid turnover and other sequence differences between PR8 and pH1N1 PA-X are responsible for their different turnover rates. Together, these data show that the region contributing PA-X instability is located between the 17^th^ and 41^st^ aa.

### PA-X amino acid 220 plays a role in protein turnover

Since sequences between PA-X aa 208 and aa 232 (X-ORF aa 17-41) appeared to control PA-X stability, we performed an alignment of the X-ORF of PA-X from the strains we tested. Although there were several amino acids that varied between the sequences, only one amino acid changed in a way that was consistent with the protein half-life, aa 220. In the longer-lived PA-X variants (Perth and pH1N1) this residue is a histidine, whereas in the shorter-lived PA-X variants (PR8, Udorn, HK06), this residue is an arginine (Fig 5A). Interestingly, this is one of only eight amino acid differences between the Udorn and Perth PA-X, which come from H3N2 strains collected almost 40 years apart and have very different half-lives in our assays (Fig 3). To determine if aa 220 plays a role in PA-X turnover, we introduced an R220H mutation in PR8 PA-X and a H220R mutation in pH1N1 PA-X. Upon a 2-hour treatment with cycloheximide, the protein levels of wt PR8 PA-X significantly decreased, but levels of PR8 PA-X R220H remained stable (Fig 5B and 5C). Conversely, levels of wt pH1N1 PA-X remained stable, but the levels of pH1N1 PA-X H220R decreased, although the change did not reach statistical significance (Fig 5E and 5F). As expected, there was no significant decrease in the mRNA levels of any of the PA-X variants in cells treated with cycloheximide (Fig 5D and 5G). Together these data point to a role for aa 220 in controlling PA-X turnover.

**Figure 5:**
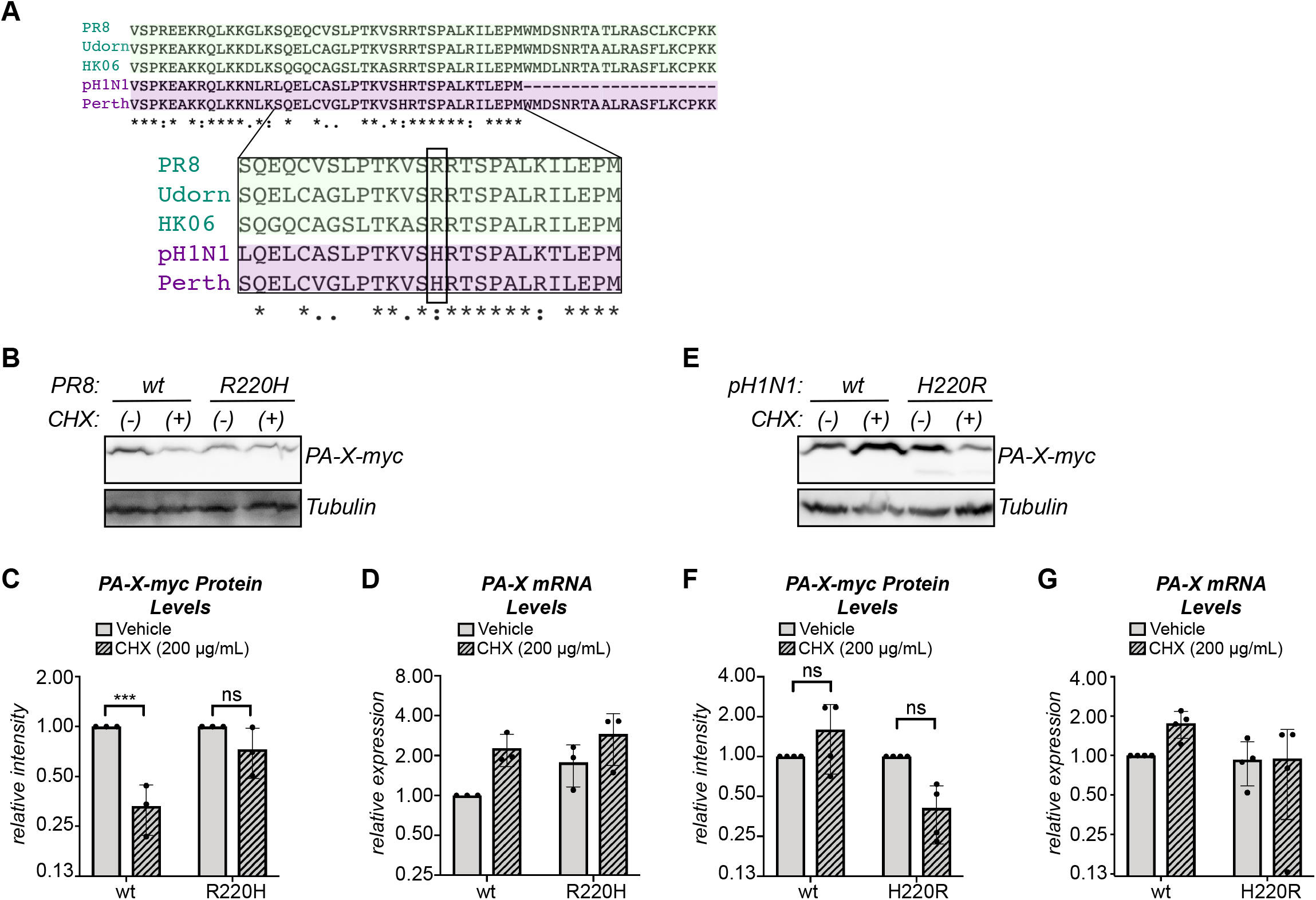
PA-X aa 220 plays a role in protein turnover. (A) Alignment of the X-ORF of PA-X from PR8, Udorn, HK06, Perth, and pH1N1. The zoomed in region shows aa 17-41 of the X-ORF (PA-X aa 208-232). Residue 220 is boxed. * = same residue, : = strong similarity,. = weak similarity. (B-G) Protein and RNA samples were collected from HEK293T cells expressing the indicated catalytically inactive C-terminally myc-tagged PA-X variant 2 hours after treatment with 200 μg/mL of cycloheximide (CHX) to inhibit translation (+) or vehicle control (−). (B, E) Representative western blot using anti-myc antibodies to detect myc-tagged PA-X with tubulin as a loading control. (C, F) For each replicate, PA-X protein levels were normalized to tubulin protein levels and reported as the amount normalized to vehicle control for each variant. *** = *p <* 0.001, ns = *p >* 0.05, 2-way ANOVA followed by Sidak’s multiple comparisons test. n ≥ 3. (D, G) Levels of PA-X mRNA were measured by RT-qPCR. After normalization to 18S rRNA, levels are plotted as amounts normalized to vehicle control treated cells expressing the wt variant. n ≥ 3.

### The N-terminal domain of PA-X also has a role in PA-X stability

To further examine how the different X-ORF variants of PR8 and pH1N1 affect PA-X stability, we created three different PR8 PA-X/pH1N1 PA-X chimeras and tested their stabilities. We extended the X-ORF of pH1N1 PA-X to 61 aa with the 20 distal residues of the PR8 X-ORF (pH1N1-X61), and we also swapped the X-ORFs of PR8 PA-X and pH1N1 PA-X (Fig 6A). Addition of the 20 distal residues of PR8 did not reduce pH1N1 stability, as pH1N1-X61 protein levels remained stable after 2 hours of cycloheximide treatment (Fig 6B and 6C), confirming that the 20 distal residues do not drive rapid turnover of PR8 PA-X. However, addition of the entire PR8 X-ORF to the pH1N1 RNase domain (pH1N1-PR8) also did not reduce the stability of the protein to the levels of PR8 PA-X after 2 hours of cycloheximide treatment (Fig 6E and 6F). In both cases, we saw a similar pattern of stability using a 4-hour cycloheximide treatment (data not shown). Furthermore, addition of the pH1N1 X-ORF to the PR8 RNase domain (PR8-pH1N1) did not increase its stability, as its protein levels still significantly decreased after 2 hours of cycloheximide (Fig 6E and 6F). In all cases there were no significant decreases in PA-X mRNA between vehicle- and cycloheximide-treated cells, confirming that any decrease in protein levels is not due to changes at the transcript level (Fig 6D and 6G). While these data are consistent with the idea that the 20-aa truncation is not responsible for the higher stability of pH1N1 PA-X, they also suggest that X-ORF sequences alone do not explain all the differences in turnover between PR8 and pH1N1 PA-X. We conclude that both the X-ORF and the N-terminal RNase domain have a role in controlling PA-X turnover rates.

**Figure 6:**
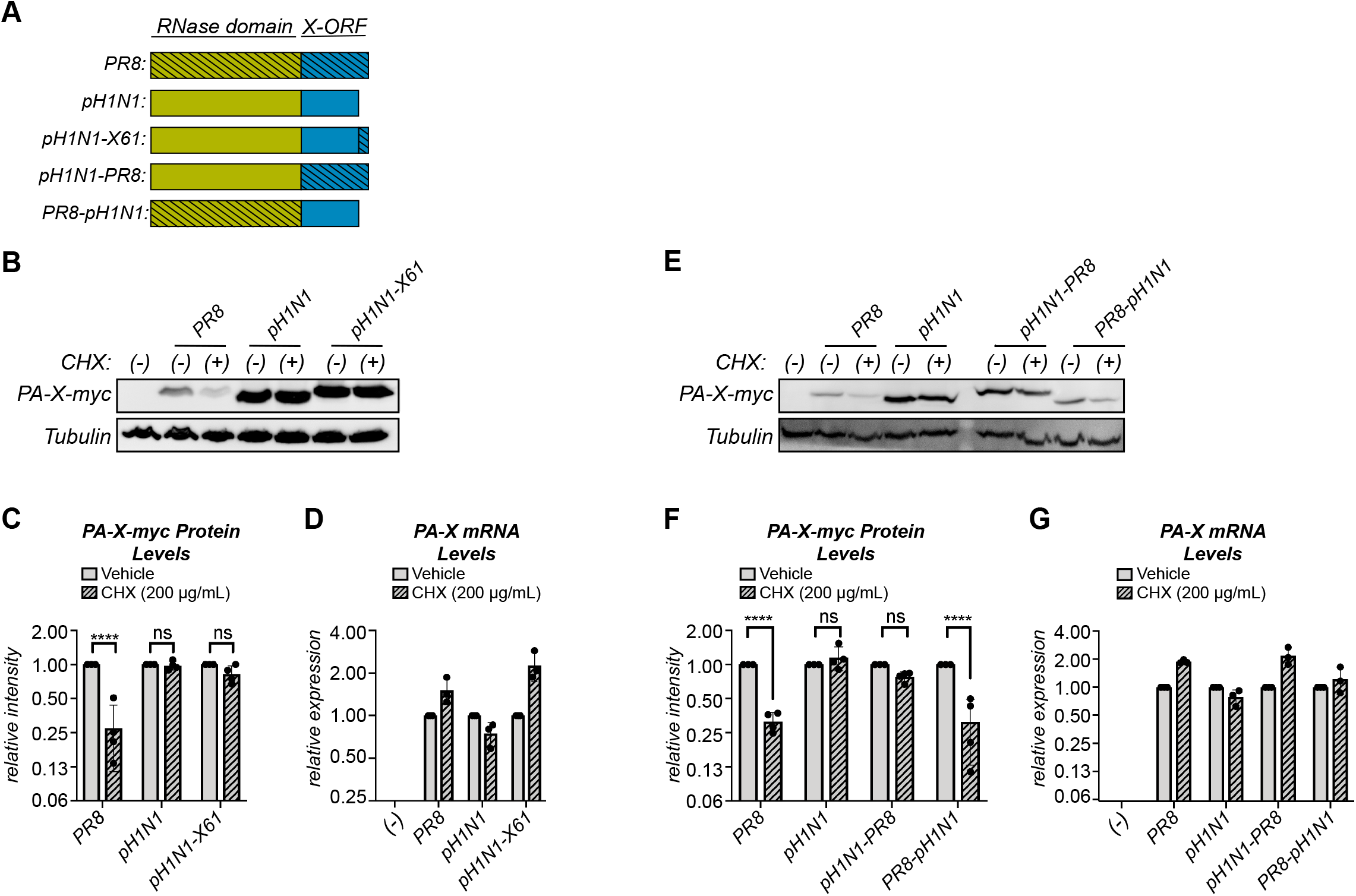
The N-terminal domain of PA-X also has a role in PA-X stability. (A) Schematic diagram of the PR8/pH1N1 PA-X chimeras used. (B-G) Protein and RNA samples were collected from HEK293T cells expressing the indicated catalytically inactive C-terminally myc-tagged PA-X variant 2 hours after treatment with 200 μg/mL of cycloheximide (CHX) to inhibit translation (+) or vehicle control (−). (B, E) Representative western blot using anti-myc antibodies to detect myc-tagged PA-X and tubulin as a loading control. (C, F) For each replicate, PA-X protein levels were normalized to tubulin protein levels and reported as the amount normalized to vehicle control for each variant. n = 4 (D, G) Levels of PA-X mRNA were measured by RT-qPCR. After normalization to 18S rRNA, levels are plotted as amounts relative to vehicle-treated cells for each variant. n = 3. **** = *p* < 0.0001, ns = *p* > 0.05, 2-way ANOVA followed by Sidak’s multiple comparisons test.

## DISCUSSION

The PA-X protein is present at low levels in influenza A virus-infected cells, which has been generally assumed to be due to its production mechanism based on ribosomal frameshifting. However, here we report that PA-X protein levels are also regulated at the post-translational level, and that PA-X is rapidly turned over (Fig 1-3). Results for host shutoff activity in PR8-infected cells suggest that PR8 PA-X is also unstable during infection and that the short half-life is not an artifact of ectopic expression (Fig 2). Moreover, protein instability is a conserved property of PA-X, because all of the PA-X variants we tested from different influenza A viruses have short half-lives, ranging from ∼30 minutes (Udorn PA-X) to ∼3.5 hours (pH1N1 PA-X) (Fig 3). At least some of the determinants of protein stability are located in the X-ORF of the protein, because truncation of the X-ORF to 16 aa significantly increases stability and fusion to the X-ORF is sufficient to induce turnover of an unrelated protein (Fig 4). Moreover, sequences between PA-X aa 208-232 (aa 17-41 of the X-ORF) and aa 220 (aa 29 of the X-ORF) in particular appear to be crucial for PA-X turnover. Truncating PA-X at aa 232 does not change its half-life (Fig 4J), while truncating it at aa 207 significantly increases it (Fig 4F), and switching aa 220 between a histidine and an arginine changes PA-X stability (Fig 5). However, the N-terminal RNase domain also has a role in PA-X turnover, as swapping the X-ORFs of PR8/pH1N1 is not sufficient to alter their stability (Fig 6). We propose that the purpose of this post-translational regulation of PA-X is to maintain low steady-state levels of the protein in infected cells, which in turn may limit the active pool of PA-X. Given that PA-X is already made at low levels during infection, the instability of this protein suggests that limiting PA-X activity is of paramount importance for viral replication.

It is attractive to think that the rapid turnover of active PA-X in the cell could serve as a mechanism to control PA-X host shutoff activity. Prolonged PA-X activity could in theory have catastrophic consequences not only for the cell but also for successful viral infection. We have observed that prolonged PA-X expression kills human cells (data not shown), as reported in yeast (13, 14). This cytopathic effect could be due to the activation of cellular stress response pathways as a response to prolonged widespread RNA degradation, or to a direct effect of the RNA degradation on the levels of unstable proteins that keep the cell alive, like the Bcl-2 family member MCL-1 (34). During infection, this toxicity could prematurely kill the infected cell before progeny virions are made, which would curtail viral replication. In support of the idea that rapid turnover limits PA-X host shutoff activity, the Takimoto group showed that truncation of the pH1N1 PA-X X-ORF to 15 residues in the virus through a premature stop codon increased host shutoff during infection (17). Based on our results, this truncation will also make PA-X more stable. Thus, the link between PA-X stability, host shutoff activity, and virulence should be further studied and rigorously tested during infection.

Several lines of evidence suggest that PA-X protein turnover is primarily regulated by the X-ORF, including changes in PA-X stability upon X-ORF truncation and aa 220 mutations and destabilization of DsRed by fusion to the X-ORF (Figs 4-5). However, the experiments with chimeric pH1N1 PA-X/PR8 PA-X also suggests a role for the N-terminal RNase domain in regulating PA-X turnover rates (Fig 6). In order to tease apart of the specific contributions of each domain to PA-X turnover rates, a more thorough understanding of the mechanism of PA-X turnover is needed. In a proteomic screen, we identified three components of the proteasome, PSME3, PSMD3, and PSMC5, and the ubiquitin E3 ligase HUWE1 as high-confidence PA-X interacting partners (3). These observations suggest a potential role for proteasomal degradation in PA-X turnover. Determining the mechanism of PA-X turnover in the infected cells and the potential role of the ubiquitin-proteasome system in this process is a current avenue of investigation.

Since the first 15 amino acids of the X-ORF are sufficient for host shutoff activity (3, 17, 22), why the rest of the X-ORF is retained through evolution has remained unclear. With the exception of a few equine H7N7 and bat influenza strains, all influenza A strains have PA-X with X-ORFs of 41 or 61 amino acids, even though there are earlier positions in the X-ORF where a stop codon could have evolved without changing the PA reading frame (16). Our study suggests that the rest of the X-ORF may be needed to regulate PA-X, and in particular to ensure that PA-X is rapidly turned over. A post-translational mechanism to control PA-X protein levels may be important to limit activity, because in the context of infection PA-X does not cleave its own transcript and thus does not autoregulate its expression (5). This is in contrast to other viral host shutoff RNases like the alphaherpesvirus protein vhs, which is thought to be able to target its own transcript (35). If hyperactive PA-X is detrimental to viral fitness, there may be selective pressure to maintain the instability of PA-X, and thus retain the distal portion of the X-ORF even though it is not required for host shutoff activity.

It was surprising that the turnover rate of pH1N1 PA-X was dramatically different from that of most other strains (Fig 3). While the biggest difference between pH1N1 and the other tested strains is the absence of the 20 distal amino acids, we find that this truncation is not responsible for the reduced degradation (Fig 4J and 6B-C). PA-X from strains with avian origin PA segments, like that of pH1N1 (36), have been reported to have higher host shutoff activity compared to PA-X from strains with human-adapted PA segments (23). This has raised the possibility that PA-X host shutoff activity is linked to virulence and pathogenicity (37). Given that pH1N1 PA-X accumulates to higher levels in the cell when compared to its counterparts from PR8, Udorn, and HK06 (Fig 1 and 3), which are human-adapted, it is possible that strain-specific differences in PA-X activity may be the result of strain-specific differences in stability. Interestingly, four amino acid changes have arisen in pH1N1 PA-X since after the 2009 pH1N1 strain was introduced into humans (24). These changes appear to reduce PA-X host shutoff activity (24). It would be interesting to investigate whether they also affect PA-X stability. In general, it will be important to assess the functional consequences of PA-X stability in terms of PA-X host shutoff activity and viral pathogenesis.

Collectively our results reveal for the first time that PA-X is markedly unstable. The stability of different PA-X variants may control host shutoff activity, which could have consequences for virulence and pathogenesis. Interestingly, we found that aa 17-41 of the X-ORF contribute to regulation of PA-X stability. Previous sequence analyses have shown that this region is maintained through evolution despite not being required for host shutoff activity. Thus, having an unstable PA-X may be advantageous for the virus and be selected for during evolution. Further elucidation of the mechanism of PA-X turnover in the infected cell and its functional consequences will uncover important aspects of PA-X biology and regulation of host shutoff, allowing us to further link PA-X activity with influenza A virus virulence and pathogenesis.

## ACKNOWLEDGEMENTS

We thank Dr. Seema Lakdawala, Dr. Chris Sullivan, Dr. Jesse Bloom and Dr. Richard Webby for constructs, and members of the Moore and Bohm lab and Dr. Malavika Raman for suggestions and feedback. We also thank Dr. Denys Khaperskyy, Dr. Craig McCormick and members of the Gaglia lab for suggestions, feedback, and critical reading of the manuscript. This work was supported by NIH grant R01 AI137358 to M.M.G. R.E.L. was supported by NIH training grant T32 AI007422. S.D.S. is a David J. Cosman, Ph.D. fellow in the Tufts Graduate School of Biomedical Sciences Molecular Microbiology Ph.D. program.

## MATERIALS AND METHODS

### Plasmids

pCR3.1-PR8-PA-X-D108A-myc was previously described (5, 38). pCR3.1-PA-X D108A-myc constructs for A/California/7/2009pdm, A/Udorn/307/1972, and A/Hong Kong/218847/2006 were generated from the corresponding previously described wild-type plasmids (3, 5) by site-directed mutagenesis. The indicated PA-X coding regions were PCR-amplified using primers in Table 2 and inserted into the SalI and XhoI sites of a pCR3.1-C-terminal-myc backbone. pCR3.1-PA-X-D108A-myc constructs for A/Perth/16/2009 (Perth) was generated by subcloning the PA-X coding region from the plasmid from segment 3 for this strain (a kind gift from Dr. Seema Lakdawala). The Perth PA-X coding region was PCR-amplified using primers in Table 2 to delete a base (which mimics the frameshift) and insert the D108A mutation. It was then inserted into the SalI and MluI sites of a pCR3.1-C-terminal-myc backbone. pCR3.1-PR8-PA-X16-D108A-myc and pCR3.1-PR8-PA-X41-D108A-myc were generated by PCR-amplifying the truncated coding region from pCR3.1-PA-X-PR8-D108A-myc using primers in Table 2. The truncated regions were then inserted into the SalI and MluI sites of a pCR3.1-C-terminal-myc backbone. pCR3.1-PR8-PA-X-D108A-R220H-myc and pCR3.1-pH1N1-PA-X-D108A-H220R-myc were generated from the corresponding D108A plasmids by site-directed mutagenesis. The indicated PA-X coding regions were PCR amplified using primers in Table 2 and inserted into the SalI and XhoI sites of pCR3.1-C-terminal-myc backbone. pCR3.1-pH1N1-X61-PA-X-D108A-myc, pCR3.1-pH1N1-PR8-PA-X-D108A-myc, pCR3.1-PR8-pH1N1-PA-X-D108A-myc were generated by amplifying the coding regions from the corresponding D108A plasmids using primers in Table 2 and then inserting the coding sequences into the SalI and XhoI sites of a pCR3.1-C-terminal-myc backbone. pCDNA3.1-dsRed-XORF and pCDNA3.1-dsRed were generated by amplifying the coding regions from the corresponding regions from pCR3.1-PR8-PA-X and pcDNA3.1-dsRed-DR (a kind gift of Chris Sullivan). The regions were then inserted into the AgeI and XhoI site of pCDNA3.1. pTRIPZ-PR8-PA-X-myc-D108A was previously described in (3, 5). Gibson cloning using HiFi assembly mix (New England Biolabs) was used to make all constructs.

**TABLE 2:**
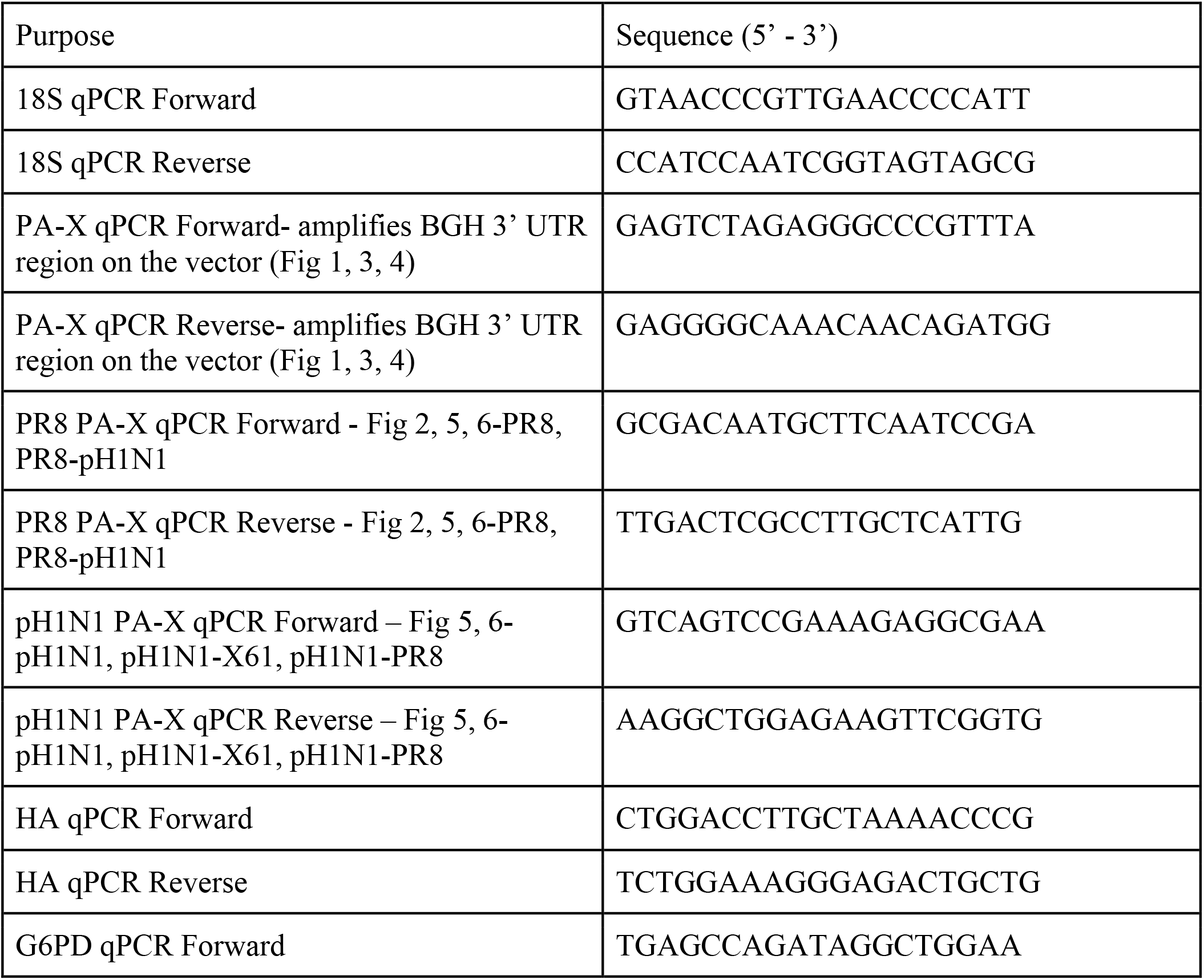
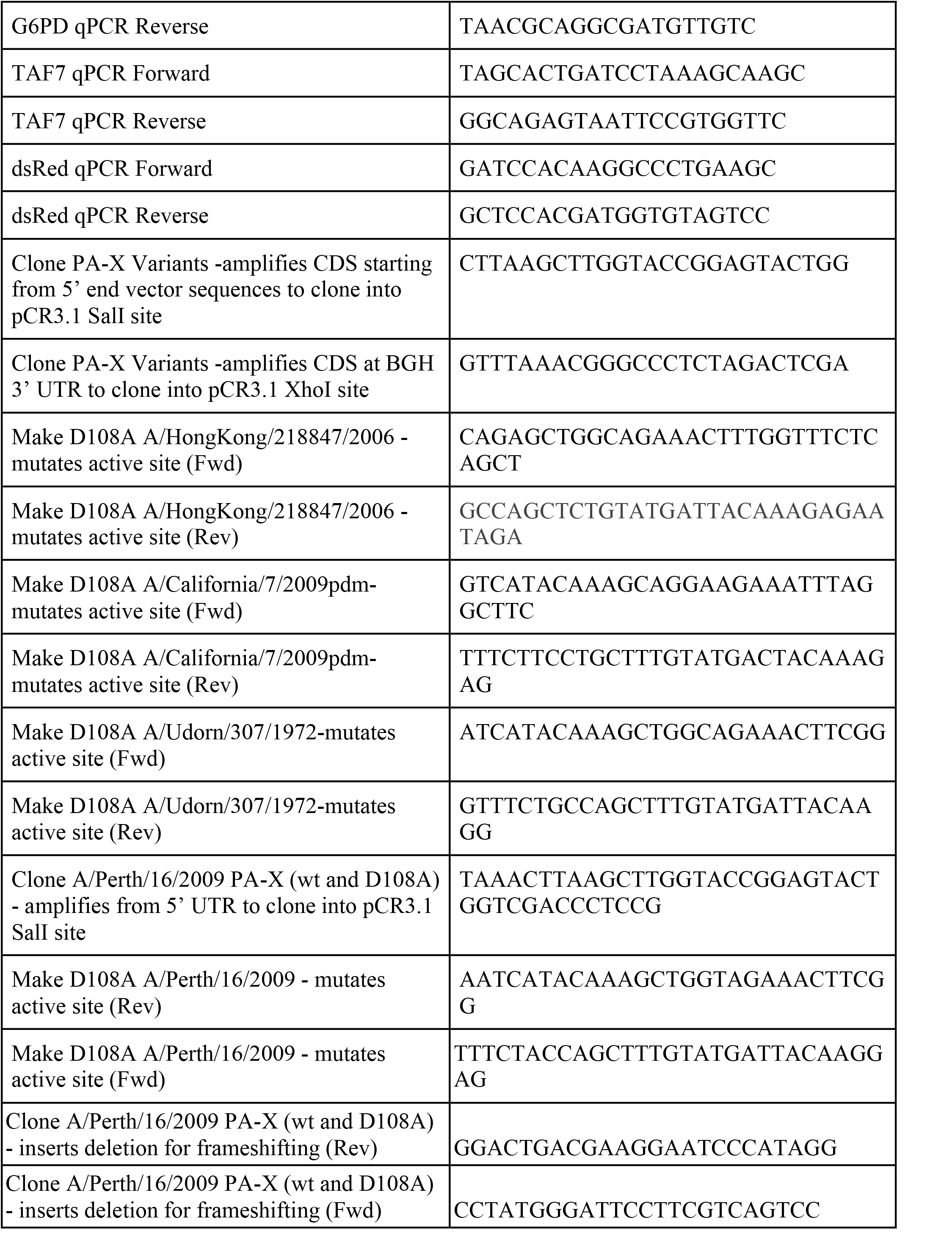
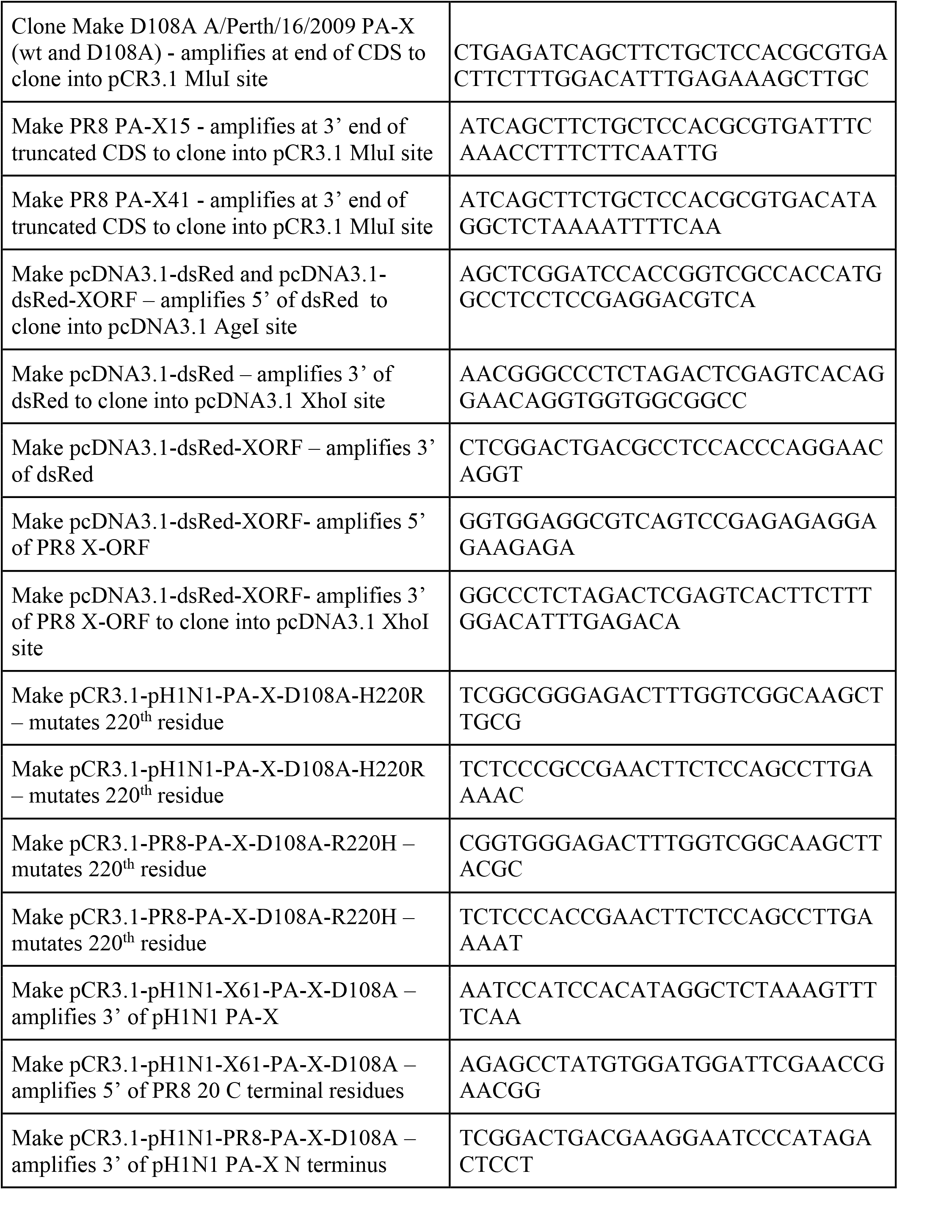
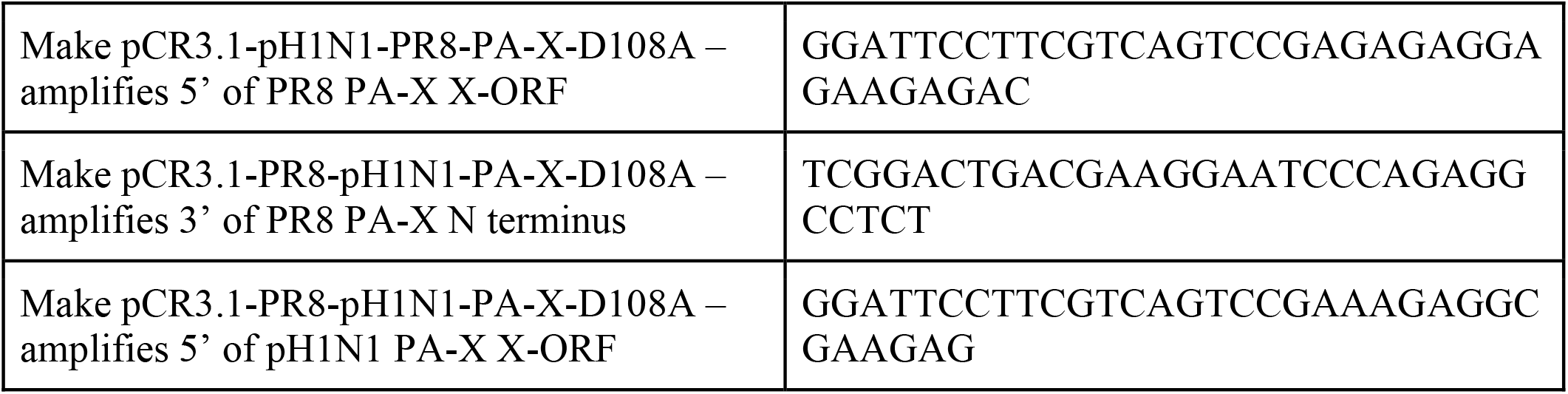
PRIMERS USED IN THIS STUDY. Primers used in this study for RT-qPCR and cloning.

### Cell lines, transfections, and drug treatments

Human embryonic kidney (HEK293T) cells (ATCC) and derivatives, Madin-Darby Canine Kidney (MDCK) cells (ATCC), A549 (ATCC) cells and derivatives were cultured in Dulbecco’s modified Eagle’s medium (DMEM; Life Technologies) supplemented with 10% fetal bovine serum (Hyclone) and maintained at 37°C in 5% CO_2_ atmosphere. A549 cell lines expressing PA-X under a doxycycline-inducible promoter (A549-iPA-X_PR8_wt#1 and A549-iPA-X_PR8_D108A#8) were previously described (5). HEK 293T-iPA-X_PR8_D108A were generated by lentiviral transduction using pTRIPZ-PR8-PA-X-myc-D108A. For all half-life experiments, HEK293T cells were plated in 10-cm dishes and transfected with 12 μg total DNA (including 6 μg of PA-X construct) using JetPrime (Polyplus transfection). 6 hours post transfection, cells were transferred to 6-well plates, and one well was set up per indicated time point. For the indicated 2, 4, and 6-hour cycloheximide experiments, HEK293T cells were plated in 2 6-wells for each indicated strain/construct and transfected with 2 μg of DNA (including 1 μg of PA-X/dsRed construct) per well using JetPrime (Polyplus transfection). 6 hours post transfection, the 2 wells for each construct were trypsinized, mixed together, and replated into 2 wells. 24 hours post transfection, cells were treated with 200 μg/mL of cycloheximide (Sigma Aldrich) or dH_2_O as vehicle control, and collected for RNA and protein isolation at the indicated time points. For experiments using A549 iPA-X cells, cells were treated with 0.5 μg/mL doxycycline (Thermo Fisher) for 14 hours to induce PA-X expression and then 200 μg/mL of cycloheximide (or vehicle control, dH_2_O) for 2 hours prior to RNA collection. For experiments using HEK293T iPA-X cells, cells were treated with 2.0 μg/mL doxycycline (Thermo Fisher) for 6 hours to induce PA-X expression prior to infection and then 200 μg/mL of cycloheximide (or vehicle control, dH_2_O) for 2 hours prior to protein collection.

### Viruses and Infections

Wild type influenza A virus A/Puerto Rico/8/1934 H1N1 (PR8) and the mutant PR8 ΔX were generated as previously described in (3) using the 8-plasmid reverse genetic system (39). Viral stocks were produced in MDCK cells and infectious titers were determined by plaque assay in MDCKs using a 1.2% Avicel overlay described in Matrosovich *et al*. (40). Cell monolayers were mock infected or infected with the wild-type or PA-X mutant viruses for 1 hr at 37°C. Monolayers were then washed with PBS, and fresh infection media (0.5% Bovine Serum Albumin (BSA; Sigma Aldrich) in DMEM was added and cells were incubated at 37°C in 5% CO_2_ atmosphere. For experiments with A549 cells, infection media was supplemented with 0.5 μg/mL of L-(tosylamido-2-phenyl) ethyl chloromethyl ketone (TPCK)-treated trypsin) (Sigma Aldrich). For A549 infections, an MOI of 5 was used, and cycloheximide was added to block translation 24 hours after infection. For HEK 293T-iPA-X_PR8_D108A infections, an MOI of 1 was used and cycloheximide was added 18 hours after infection.

### RNA purification, cDNA preparation and RT-qPCR

RNA was extracted from cells and purified using the Quick-RNA Miniprep kit (Zymo Research), following manufacturer’s protocol. In all cases, the RNA was treated with Turbo DNase (Life Technologies), then reverse transcribed using iScript supermix (Bio-Rad) per manufacturer’s protocol. qPCR was performed using iTaq Universal SYBR Green supermix (Bio-Rad), on the Bio-Rad CFX Connect Real-Time System qPCR and analyzed with Bio-Rad CFX Manager 3.1 Program. The primers used are listed in Table 2.

### Protein extraction, western blotting and antibodies

Total cellular protein was collected in protein lysis buffer (50 mM Tris-HCl pH 7.4, 150 mM NaCl, 2 mM EDTA, 0.5% sodium deoxycholate, 0.5% SDS, 1% NP-40) supplemented with cOmplete EDTA-free protease inhibitor cocktail (Roche) and 1 U/mL benzonase nuclease (Sigma Aldrich). The benzonase is included to degrade DNA and promote release of PA-X from the nucleus. Proteins were separated on 12% SDS-PAGE gels and transferred onto 0.22 μm PVDF membranes (Millipore). Western Blots were performed in PBST with 0.5% milk using antibodies against the Myc tag (1:1000, Cell Signaling Technologies 2276), influenza A virus (1:2000, Abcam ab20841), dsRed (1:200, Santa Cruz Biotechnology sc-101526), human GAPDH (1:1000, Cell Signaling Technologies 5174), and human β-tubulin (1:1000, Cell Signaling Technologies 2128). Secondary antibodies conjugated to HRP were purchased from Southern Biotechnologies (rabbit, OB4030-05; mouse, OB1031-05; goat, OB6106-05) and used at 1:5,000 dilution. Membranes were developed using Pierce ECL Western blotting substrate (Thermo Fisher) and imaged with a Syngene G:Box Chemi XT4 gel doc system. Images were quantified using GeneTools version 4.3.8.0

### Statistical Analysis

All statistical analysis was performed using GraphPad Prism version 8.0 for Mac OS X, (GraphPad Software, San Diego, CA, USA). For half-life experiments, statistical significance was determined by performing linear regression and comparing slopes and intercepts. For other experiments, statistical significance was determined by Student’s *t*-test, or analysis of variance (ANOVA) followed by a *post-hoc* multiple comparisons test (Sidak’s or Tukey’s) as indicated. All bar graphs are mean +/- standard deviation. A minimum of three biological replicates were conducted for all experiments.

## References

1. Jagger BW, Wise HM, Kash JC, Walters K-A, Wills NM, Xiao Y-L, Dunfee RL, Schwartzman LM, Ozinsky A, Bell GL, Dalton RM, Lo A, Efstathiou S, Atkins JF, Firth AE, Taubenberger JK, Digard P. 2012. An Overlapping Protein-Coding Region in Influenza A Virus Segment 3 Modulates the Host Response. Science 337:199–204.

2. Levene RE, Gaglia MM. 2018. Host Shutoff in Influenza A Virus: Many Means to an End. Viruses 10:475.

3. Gaucherand L, Porter BK, Levene RE, Price EL, Schmaling SK, Rycroft CH, Kevorkian Y, McCormick C, Khaperskyy DA, Gaglia MM. 2019. The Influenza A Virus Endoribonuclease PA-X Usurps Host mRNA Processing Machinery to Limit Host Gene Expression. Cell Reports 27:776–792.e7.

4. Chaimayo C, Dunagan M, Hayashi T, Santoso N, Takimoto T. Specificity and functional interplay between influenza virus PA-X and NS1 shutoff activity 26.

5. Khaperskyy DA, Schmaling S, Larkins-Ford J, McCormick C, Gaglia MM. 2016. Selective Degradation of Host RNA Polymerase II Transcripts by Influenza A Virus PA-X Host Shutoff Protein. PLOS Pathogens 12:e1005427.

6. Gao H, Sun Y, Hu J, Qi L, Wang J, Xiong X, Wang Y, He Q, Lin Y, Kong W, Seng L-G, Sun H, Pu J, Chang K-C, Liu X, Liu J. 2015. The contribution of PA-X to the virulence of pandemic 2009 H1N1 and highly pathogenic H5N1 avian influenza viruses. Scientific Reports 5:8262.

7. Gong X-Q, Sun Y-F, Ruan B-Y, Liu X-M, Wang Q, Yang H-M, Wang S-Y, Zhang P, Wang X-H, Shan T-L, Tong W, Zhou Y-J, Li G-X, Zheng H, Tong G-Z, Yu H. 2017. PA-X protein decreases replication and pathogenicity of swine influenza virus in cultured cells and mouse models. Veterinary Microbiology 205:66–70.

8. Hayashi T, MacDonald LA, Takimoto T. 2015. Influenza A Virus Protein PA-X Contributes to Viral Growth and Suppression of the Host Antiviral and Immune Responses. Journal of Virology 89:6442–6452.

9. Hu J, Mo Y, Wang X, Gu M, Hu Z, Zhong L, Wu Q, Hao X, Hu S, Liu W, Liu H, Liu X, Liu X. 2015. PA-X Decreases the Pathogenicity of Highly Pathogenic H5N1 Influenza A Virus in Avian Species by Inhibiting Virus Replication and Host Response. Journal of Virology 89:4126–4142.

10. Xu G, Zhang X, Liu Q, Bing G, Hu Z, Sun H, Xiong X, Jiang M, He Q, Wang Y, Pu J, Guo X, Yang H, Liu J, Sun Y. 2017. PA-X protein contributes to virulence of triple-reassortant H1N2 influenza virus by suppressing early immune responses in swine. Virology 508:45– 53.

11. Sun Y, Hu Z, Zhang X, Chen M, Wang Z, Xu G, Bi Y, Tong Q, Wang M, Sun H, Pu J, Iqbal M, Liu J. 2020. R195K mutation in the PA-X protein increases the virulence and transmission of influenza A virus in mammalian hosts. J Virol JVI.01817-19, jvi;JVI.01817-19v1.

12. Liu L, Song S, Shen Y, Ma C, Wang T, Tong Q, Sun H, Pu J, Iqbal M, Liu J, Sun Y. 2020. Truncation of PA-X Contributes to Virulence and Transmission of H3N8 and H3N2 Canine Influenza Viruses in Dogs. J Virol 94:e00949–20, /jvi/94/15/JVI.00949-20.atom.

13. Oishi K, Yamayoshi S, Kozuka-Hata H, Oyama M, Kawaoka Y. 2018. N-Terminal Acetylation by NatB Is Required for the Shutoff Activity of Influenza A Virus PA-X. Cell Reports 24:851–860.

14. Oishi K, Yamayoshi S, Kawaoka Y. 2018. Identification of novel amino acid residues of influenza virus PA-X that are important for PA-X shutoff activity by using yeast. Virology 516:71–75.

15. Firth AE, Jagger BW, Wise HM, Nelson CC, Parsawar K, Wills NM, Napthine S, Taubenberger JK, Digard P, Atkins JF. 2012. Ribosomal frameshifting used in influenza A virus expression occurs within the sequence UCC_UUU_CGU and is in the +1 direction. Open Biology 2:120109.

16. Shi M, Jagger BW, Wise HM, Digard P, Holmes EC, Taubenberger JK. 2012. Evolutionary Conservation of the PA-X Open Reading Frame in Segment 3 of Influenza A Virus. Journal of Virology 86:12411–12413.

17. Hayashi T, Chaimayo C, McGuinness J, Takimoto T. 2016. Critical Role of the PA-X C-Terminal Domain of Influenza A Virus in Its Subcellular Localization and Shutoff Activity. J Virol 90:7131–7141.

18. Košík I, Práznovská M,Košíková M, Bobišová Z, Hollý J, Varečková E, Kostolanský F, Russ G. 2015. The Ubiquitination of the Influenza A Virus PB1-F2 Protein Is Crucial for Its Biological Function. PLoS ONE 10:e0118477.

19. Varga ZT, Palese P. 2011. The influenza A virus protein PB1-F2. Virulence 2:542–546.

20. Varga ZT, Ramos I, Hai R, Schmolke M, García-Sastre A, Fernandez-Sesma A, Palese P. 2011. The Influenza Virus Protein PB1-F2 Inhibits the Induction of Type I Interferon at the Level of the MAVS Adaptor Protein. PLoS Pathog 7:e1002067.

21. Varga ZT, Grant A, Manicassamy B, Palese P. 2012. Influenza Virus Protein PB1-F2 Inhibits the Induction of Type I Interferon by Binding to MAVS and Decreasing Mitochondrial Membrane Potential. J Virol 86:8359–8366.

22. Oishi K, Yamayoshi S, Kawaoka Y. 2015. Mapping of a Region of the PA-X Protein of Influenza A Virus That Is Important for Its Shutoff Activity. Journal of Virology 89:8661– 8665.

23. Desmet EA, Bussey KA, Stone R, Takimoto T. 2013. Identification of the N-Terminal Domain of the Influenza Virus PA Responsible for the Suppression of Host Protein Synthesis. Journal of Virology 87:3108–3118.

24. Nogales A, Martinez-Sobrido L, Chiem K, Topham DJ, DeDiego ML. 2018. Functional Evolution of the 2009 Pandemic H1N1 Influenza Virus NS1 and PA in Humans. Journal of Virology 92:e01206–18.

25. Santos A, Pal S, Chacón J, Meraz K, Gonzalez J, Prieto K, Rosas-Acosta G. 2013. SUMOylation Affects the Interferon Blocking Activity of the Influenza A Nonstructural Protein NS1 without Affecting Its Stability or Cellular Localization. Journal of Virology 87:5602–5620.

26. Cheng Y-Y, Yang S-R, Wang Y-T, Lin Y-H, Chen C-J. 2017. Amino Acid Residues 68–71 Contribute to Influenza A Virus PB1-F2 Protein Stability and Functions. Front Microbiol 8:692.

27. Chen W M. Smeekens J, Wu R. 2016. Systematic study of the dynamics and half-lives of newly synthesized proteins in human cells. Chemical Science 7:1393–1400.

28. Rigby RE, Wise HM, Smith N, Digard P, Rehwinkel J. 2019. PA-X antagonises MAVS-dependent accumulation of early type I interferon messenger RNAs during influenza A virus infection. Sci Rep 9:7216.

29. Hara K, Schmidt FI, Crow M, Brownlee GG. 2006. Amino Acid Residues in the N-Terminal Region of the PA Subunit of Influenza A Virus RNA Polymerase Play a Critical Role in Protein Stability, Endonuclease Activity, Cap Binding, and Virion RNA Promoter Binding. Journal of Virology 80:7789–7798.

30. Maier HJ, Kashiwagi T, Hara K, Brownlee GG. 2008. Differential role of the influenza A virus polymerase PA subunit for vRNA and cRNA promoter binding. Virology 370:194– 204.

31. Gao H, Sun H, Hu J, Qi L, Wang J, Xiong X, Wang Y, He Q, Lin Y, Kong W, Seng L-G, Pu J, Chang K-C, Liu X, Liu J, Sun Y. 2015. Twenty amino acids at the C-terminus of PA-X are associated with increased influenza A virus replication and pathogenicity. Journal of General Virology 96:2036–2049.

32. Feng KH, Sun M, Iketani S, Holmes EC, Parrish CR. 2016. Comparing the functions of equine and canine influenza H3N8 virus PA-X proteins: Suppression of reporter gene expression and modulation of global host gene expression. Virology 496:138–146.

33. Wang X-H. 2020. The role of PA-X C-terminal 20 residues of classical swine influenza virus in its replication and pathogenicity. Veterinary Microbiology 5.

34. Adams KW, Cooper GM. 2007. Rapid Turnover of Mcl-1 Couples Translation to Cell Survival and Apoptosis. J Biol Chem 282:6192–6200.

35. Elgadi MM, Hayes CE, Smiley JR. 1999. The Herpes Simplex Virus vhs Protein Induces Endoribonucleolytic Cleavage of Target RNAs in Cell Extracts. J Virol 73:7153–7164.

36. Garten RJ, Davis CT, Russell CA, Shu B, Lindstrom S, Balish A, Sessions WM, Xu X, Skepner E, Deyde V, Okomo-Adhiambo M, Gubareva L, Barnes J, Smith CB, Emery SL, Hillman MJ, Rivailler P, Smagala J, Graaf M de, Burke DF, Fouchier RAM, Pappas C, Alpuche-Aranda CM, López-Gatell H, Olivera H, López I, Myers CA, Faix D, Blair PJ, Yu C, Keene KM, Dotson PD, Boxrud D, Sambol AR, Abid SH, George KS, Bannerman T, Moore AL, Stringer DJ, Blevins P, Demmler-Harrison GJ, Ginsberg M, Kriner P, Waterman S, Smole S, Guevara HF, Belongia EA, Clark PA, Beatrice ST, Donis R, Katz J, Finelli L, Bridges CB, Shaw M, Jernigan DB, Uyeki TM, Smith DJ, Klimov AI, Cox NJ. 2009. Antigenic and Genetic Characteristics of Swine-Origin 2009 A(H1N1) Influenza Viruses Circulating in Humans. Science 325:197–201.

37. Lutz MM, Dunagan MM, Kurebayashi Y, Takimoto T. 2020. Key Role of the Influenza A Virus PA Gene Segment in the Emergence of Pandemic Viruses. 4. Viruses 12:365.

38. Khaperskyy DA, Emara MM, Johnston BP, Anderson P, Hatchette TF, McCormick C. 2014. Influenza A Virus Host Shutoff Disables Antiviral Stress-Induced Translation Arrest. PLOS Pathogens 10:e1004217.

39. Hoffmann E, Neumann G, Kawaoka Y, Hobom G, Webster RG. 2000. A DNA transfection system for generation of influenza A virus from eight plasmids. Proceedings of the National Academy of Sciences 97:6108–6113.

40. Matrosovich M, Matrosovich T, Garten W, Klenk H-D. 2006. New low-viscosity overlay medium for viral plaque assays. Virol J 3:63.

